# Simplifying Systems Genetics with QTLretrievR: An R Package for Molecular QTL Identification

**DOI:** 10.64898/2026.02.16.706200

**Authors:** Hannah B. Dewey, Selcan Aydin, Samuel J. Widmayer, Daniel A. Skelly, Daniel M. Gatti, Steven C. Munger

## Abstract

**Motivation:** Over the past two decades, integrating “-omics” scale molecular profiling with genetic mapping in diverse populations—termed “systems genetics”—has proven powerful for linking complex disease phenotypes to genetic variants and their regulatory effects. However, these combined genomic and genetic analyses remain computationally intensive and often inaccessible to researchers without a strong computational background, a problem compounded in studies with multiple tissues or thousands of molecular -omics phenotypes.

**Results:** We developed QTLretrievR, an R package designed to simplify and streamline the identification and downstream analysis of molecular quantitative trait loci in genetically diverse populations. QTLretrievR builds on widely used R packages such as *r/qtl2* for quantitative trait locus mapping and *r/intermediate* for mediation analysis, and incorporates substantial parallelization features to improve computational efficiency. Benchmarking demonstrates that QTLretrievR can reproduce results from previous quantitative trait locus studies while significantly reducing the time required for quantitative trait locus peak detection and downstream analyses. Additionally, QTLretrievR includes plotting functions to visualize results of the included downstream analyses.

**Availability:** Source code is available at https://github.com/deweyhannah/QTLretrievR.

**Contact:** Steven Munger

## 1 Introduction

Quantitative trait locus (QTL) mapping links phenotypic variation among diverse individuals to genetic variation at one or more loci across the genome. While QTL mapping was originally developed for application to individual disease or physiological phenotypes, more recently it has been used to identify the genetic determinants of population variation in molecular phenotypes (e.g., transcriptomics, proteomics), an integrative approach called “systems genetics”. At its core, QTL mapping can be broken down into three phases: data normalization and preparation; genetic mapping, significance thresholding, and peak calling; and downstream analysis of identified QTL. Each of these steps involves data cleaning and transformation of some form, and these complexities can present a barrier to entry for researchers lacking statistical or computational training.

The popular R package *r/qtl2*(1) provides a standardized method for performing QTL mapping, with functions to impute and collate sample genotype probabilities from marker genotypes, and flexible models for genetic mapping, peak calling, and estimation of genotype effects at the QTL. However, *r/qtl2* analyzes phenotypes in series, and scaling up to thousands of molecular phenotypes that are routinely measured today with -omics platforms necessitates considerable computational resources and adds significant complexity to QTL mapping and downstream analyses. Despite these challenges, the upside to QTL mapping with thousands of molecular traits is the added power to infer meaningful genetic relationships genome-wide. For example, genetic variant(s) that drive population variation in higher-order phenotypes (e.g., cell differentiation bias, organ function, disease susceptibility) may confer these consequences through direct, proximal effects on gene expression regulation and also map as a gene expression QTL (eQTL). In fact, genome wide association studies (GWAS) in human populations have consistently found that most complex trait-associated variants map to non-coding regions and operate through regulatory mechanisms, as opposed to coding variants that directly affect protein structure or function(2, 3).

A strength of systems genetics approaches is the ability to leverage genetic variation across the genome to infer causal relationships between genetic, molecular, and cellular or organismal phenotypes. Mediation analysis is a powerful tool used to identify these causal “mediator” genes where the genetic variation is influencing the expression of regulatory genes with downstream effects on hundreds of other genes leading to variation in higher-order phenotypes(4–6). Performing these analyses within *r/qtl2* requires numerous steps and complex data handling and filtering steps; alternatively, QTL mapping results from *r/qtl2* can be exported to a mediation analysis R package such as *r/intermediate*(4) or *r/bmediatR*(7). However, these stand alone software packages come with their own challenges, including a lack of parallelization (*r/intermediate*) and constraints on the numbers of phenotype-mediator pairs that can be handled at a time (*r/bmediatR*).

We developed *r/QTLretrievR* to both simplify the process and increase the computational efficiency of performing molecular QTL mapping and mediation analysis at scale by incorporating *r/qtl2* and *r/intermediate* into a single, end-to-end pipeline. By adding parallelization to take advantage of available compute resources, QTLretrievR is able to process thousands of quantitative phenotypes from multiple tissue types simultaneously. QTL peak calling and mediation analyses can be performed using a minimal number of commands, and the outputs of the mapping and peak calling steps can be ported directly to mediation analysis. In all, QTLretrievR includes functions to: prepare sample genotype probabilities; estimate “logarithm of the odds” (LOD) significance thresholds empirically with permutation testing; perform additive and interactive QTL mapping and peak calling; identify genomic loci enriched for distant QTL, i.e. QTL “hotspots”; and analyze distant QTL with mediation analysis to nominate candidate regulatory genes for later experimental validation. Importantly, results from these analyses can be visualized and output as publication-ready plots. In summary, QTLretrievR is an easy-to-use yet robust pipeline for molecular QTL mapping that is designed to support both novice and experienced computational users. Although originally developed for use with the Diversity Outbred (DO) and Collaborative Cross (CC) mouse populations, QTLretrievR is broadly applicable to any genetically diverse population with available genotype and phenotype data.

## 2 Methods

### 2.1. Overview

QTLretrievR is an R package that provides flexible functions for performing QTL analysis on high-dimensional “-omics” datasets comprising thousands of molecular phenotypes (e.g., RNA-seq, proteomics), from single or multiple types of biosamples (e.g., tissues, cell types), and hundreds of genetically diverse individuals. QTLretrievR includes data preprocessing and analysis steps, as well as data visualization functions that have been optimized to leverage available computational resources. In particular, the peak calling, genotype effect calculation, and mediation analyses are implemented with parallelization that takes the total number of features and available cores into account to maximize computational efficiency. QTLretrievR was built in R version 4.3.1, with devtools 2.4.5. It uses the native R pipe, which is not compatible with R versions < 4.1.

### 2.2. Diversity Outbred Mice

QTLretrievR was originally designed to support molecular QTL analyses using the Diversity Outbred (DO) mouse panel. The DO is a heterogeneous stock created by intercrossing eight inbred “founder” strains and maintained by random outbreeding among 175 breeding pairs (8). The eight founder strains include five common inbred laboratory strains A/J, C57BL/6J, 129S1/SvImJ, NOD/ShiLtJ, and NZO/HlLtJ, in addition to three more recently wild-derived inbred strains representing three mouse subspecies, *M. musculus domesticus* (WSB/EiJ), *M. musculus musculus* (PWK/PhJ), and *M. musculus castaneus* (CAST/EiJ). Together, these eight strains segregate 40 million single nucleotide polymorphisms (SNPs) as well as millions of short insertion/deletions (indels) and larger structural variants(9). This abundant genetic variation is broadly spread throughout the genome, and has been shown to modify the expression of thousands of genes in a tissue-specific or conserved manner(4–6, 10–13).

### 2.3. Implementation

#### 2.3.1. Parallelization

QTLretrievR introduces an additional layer of parallelization across QTL mapping and peak calling, genotype effect inference, and mediation. When there are more than 1000 phenotypes, they are split into batches and distributed over available computational cores for processing. The parallelization is implemented using the *r/doParallel*(14) package, and the number of available cores is detected by system inquiry. Multi-tissue or multi-condition analyses are supported by dividing available cores among tissues/conditions before dividing them between phenotypes. Results for each tissue are returned in a single unified object. While this improves flexibility, efficiency gains will depend on computational resources available to the end user.

#### 2.3.2. Significance Thresholding

Permutation testing is used to estimate genome-wide significance for mapped QTL. In permutation testing, the rows of the genotype data are randomly shuffled (permuted) among samples while the relationship between the phenotypes and any sample covariates (e.g., sex, treatment group) is maintained. For each permutation, the maximum LOD score is retained to generate a null distribution for the test statistic calculation(15). For comparison to established methods, we referenced our published eQTL study of mouse DO neural progenitor cells (NPCs) that performed permutation testing on every individually expressed gene(6) (hereafter referred to as “genespecific thresholding”; n = 14163 genes total). We randomly sampled autosomal genes for permutation testing to avoid sex chromosome biases, and varied the number of subsampled genes between 10 - 100 and the permutations per gene between 500 - 1000. The median significant LOD threshold for each combination was compared to the ground truth, gene-specific LOD thresholds, and accuracy 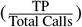, precision and recall (*r/caret*(16)) were calculated for each of 1000 simulations. Equivalence for each combination was determined using a one sample two one sided tests (TOST) on accuracy - 1 for all simulations, with an epsilon 0.03.

#### 2.3.3. Streamlining

QTLretrievR enhances reproducibility and simplifies user interaction by streamlining QTL analysis steps within the implementation. We converted existing, regularly referenced scripts that used different analysis packages into modular functions to standardize their inputs and outputs. Resources that are stable and required for multiple analysis types (e.g., SNP marker maps and gene annotations) are included as objects that are automatically loaded with the package. Extensive testing ensured that outputs generated from earlier steps were formatted to match the expected inputs of subsequent steps. Each step of the analysis returns a single list of objects, organized by tissue (if multiple tissues are analyzed), containing all elements needed for the subsequent stage.

### 2.4. Benchmarking

Benchmarking was performed to evaluate the computational efficiency and scalability of QTLretrievR relative to *r/qtl2*(1). For these analyses, we utilized a published pancreatic islet RNA-seq dataset with 22180 gene expression phenotypes from 378 DO mouse samples(11, 17–19), and adapted an *r/qtl2* script from an internal eQTL training course to test the ‘scan1’ mapping function with variable numbers of cores. The script was run five times with varying resources (described further in Table 1), keeping available memory and maximum job time consistent. A point of diminishing returns was determined by creating a linear model and using a combination of segmentation and estimated marginal means analyses. For comparison, QTLretrievR was passed the same set of total cores, with varying minimum cores passed to ‘scan1’ to test combinations of the number of parallel processes and number of cores (described further in Table 1).

**Table 1.** Time (hours) to complete mapping with and without parallelization, 5 replicates.

| Total | Cores | Procs | Rep1 | Rep2 | Rep3 | Rep4 | Rep5 |
| --- | --- | --- | --- | --- | --- | --- | --- |
| 4 | 4 | 1 | 43.32 | 42.93 | 44.96 | 43.09 | 39.67 |
| 8 | 8 | 1 | 23.87 | 25.72 | 24.44 | 24.31 | 24.01 |
| 12 | 4 | 3 | 20.5 | 21.55 | 23.18 | 21.95 | 20.32 |
| 12 | 12 | 1 | 17.15 | 17.78 | 17.62 | 17.04 | 18.19 |
| 16 | 4 | 4 | 17.52 | 15.17 | 17.74 | 17.62 | 17.51 |
| 16 | 8 | 2 | 15.83 | 16.48 | 16.12 | 16.33 | 17.32 |
| 16 | 16 | 1 | 13.48 | 13.85 | 14.06 | 13.81 | 15.37 |
| 20 | 4 | 5 | 13.75 | 14.28 | 13.89 | 12.69 | 13.74 |
| 20 | 20 | 1 | 13.14 | 13.13 | 12.79 | 13.2 | 12.94 |
| 24 | 4 | 6 | 12.73 | 11.18 | 11.46 | 11.44 | 11.42 |
| 24 | 8 | 3 | 12.07 | 11.34 | 11.13 | 11.89 | 12.14 |
| 24 | 12 | 2 | 11.01 | 10.63 | 11.92 | 11.05 | 11.17 |
| 24 | 24 | 1 | 11.45 | 12.48 | 10.45 | 10.42 | 12.35 |
| 28 | 4 | 7 | 9.78 | 9.71 | 10.59 | 12.12 | 10.89 |
| 28 | 28 | 1 | 10.18 | 10.5 | 10.21 | 9.84 | 10.35 |
| 36 | 4 | 9 | 7.22 | 7.1 | 7.35 | 7.62 | 7.05 |
| 36 | 8 | 4 | 7.18 | 7.16 | 5.25 | 5.26 | 5.29 |
| 36 | 12 | 3 | 5.1 | 8.11 | 5.11 | 5.06 | 5.07 |
| 36 | 16 | 2 | 5.19 | 5.2 | 5.22 | 5.21 | 7.38 |
| 36 | 20 | 1 | 9.01 | 9.81 | 10.34 | 10.9 | 12 |
| 36 | 24 | 1 | 9.87 | 9.97 | 9.07 | 9.56 | 8.75 |
| 36 | 36 | 1 | 7.19 | 6.72 | 6.9 | 8.61 | 7.29 |
| 44 | 4 | 11 | 6.76 | 6.48 | 7.1 | 6.07 | 6.15 |
| 44 | 8 | 5 | 6.58 | 6.56 | 6.84 | 6.47 | 6.41 |
| 44 | 12 | 3 | 6.74 | 7.67 | 6.61 | 6.56 | 6.81 |
| 44 | 16 | 2 | 7.52 | 7.81 | 7.68 | 7.91 | 7.26 |
| 44 | 20 | 2 | 8.47 | 8.35 | 8.09 | 7.57 | 7.71 |
| 44 | 24 | 1 | 8.88 | 8.22 | 8.73 | 8.76 | 8.85 |
| 44 | 44 | 1 | 6.65 | 6.77 | 6.19 | OOM | OOM |
| 52 | 4 | 13 | 5.78 | 5.59 | 5.83 | 6.17 | 5.83 |
| 52 | 8 | 6 | 5.44 | 6.53 | 5.8 | 6.28 | 5.83 |
| 52 | 12 | 4 | 5.49 | 5.96 | 5.49 | 5.76 | 6 |
| 52 | 16 | 3 | 6.04 | 5.72 | 5.84 | 5.78 | 5.9 |
| 52 | 20 | 2 | 6.72 | 6.7 | 6.43 | 7.02 | 6.73 |
| 52 | 24 | 2 | 5.52 | 5.98 | 5.89 | 5.51 | 5.88 |
| 52 | 52 | 1 | OOM | OOM | OOM | OOM | OOM |
| 60 | 4 | 15 | 6.08 | 5.35 | 5.65 | 5.72 | 5.81 |
| 60 | 8 | 7 | 5.09 | 5.83 | 5.88 | 5.69 | 5.92 |
| 60 | 12 | 5 | 4.98 | 5.14 | 5.38 | 5.51 | 5.26 |
| 60 | 16 | 3 | 5.43 | 5.34 | 5.38 | 5.38 | 5.51 |
| 60 | 20 | 3 | 5.56 | 5.86 | 6.01 | 6.26 | 5.56 |
| 60 | 24 | 2 | 5.2 | 5.34 | 5.18 | 5.22 | 5.24 |
| 60 | 60 | 1 | OOM | OOM | OOM | OOM | OOM |
| 68 | 4 | 17 | 5.27 | 5.22 | 5.16 | 5.32 | 5.12 |
| 68 | 8 | 8 | 4.26 | 4.79 | 4 | 4.28 | 4.08 |
| 68 | 12 | 5 | 4.83 | 4.85 | 4.6 | 4.72 | 4.88 |
| 68 | 16 | 4 | 4.1 | 4.61 | 4.11 | 4.2 | 4.55 |
| 68 | 20 | 3 | 5.64 | 5.64 | 5.55 | 5.38 | 5.45 |
| 68 | 24 | 2 | 5.37 | 5.19 | 4.37 | 4.83 | 4.41 |
| 68 | 68 | 1 | OOM | OOM | OOM | OOM | OOM |
| 72 | 4 | 18 | 4.44 | 4.34 | 4.36 | 4.58 | 4.37 |
| 72 | 8 | 9 | 4.26 | 4.24 | 3.95 | 3.92 | 4.44 |
| 72 | 12 | 6 | 4.57 | 4.54 | 4.59 | 3.74 | 3.74 |
| 72 | 16 | 4 | 3.96 | 3.9 | 3.95 | 4.24 | 4.01 |
| 72 | 20 | 3 | 4.47 | 5.17 | 5.32 | 5.24 | 5.32 |
| 72 | 24 | 3 | 3.76 | 3.76 | 3.8 | 3.75 | 3.74 |
| 72 | 72 | 1 | OOM | OOM | OOM | OOM | OOM |

## 3 Results and Discussion

### 3.1. Mapping

To optimize the efficiency of QTLretrievR for large-scale molecular QTL analyses, we determined the saturation point at which passing additional cores to *r/qtl2*(1) no longer significantly decreased the amount of time to run mapping for thousands of phenotypes. For this analysis, we used the pancreatic islet RNA-seq dataset with 22180 expressed genes from 378 DO mouse samples(11). We found that increasing the total number of cores passed to *r/qtl2* decreased mapping time, but efficiency gains diminished beyond 12 cores. Segmented regression identified a breakpoint between 8 and 12 cores, and pairwise comparisons confirmed significant improvements up to 16 cores (BH adj. p < 0.0001). Significant differences were also found between 28 and 36 cores, but that is due to the step between cores switching from 4 to 8. There were no significant differences for other allocations (BH adj. p > 0.01).

To assess how different configurations of cores allocated to ‘scan1’ and different numbers of parallel processes affected efficiency, we passed QTLretrievR between 12 and 72 total cores and set the minimum number of cores used per mapping process to be less than the total number of cores (Table 1). For total cores <24, the addition of parallelization did not reduce the amount of time needed to complete mapping. For 24 total cores, the average time required to complete each parallelization configuration (4 cores x 6 processes, 8 cores x 3 processes, 12 cores x 2 processes) matched or slightly reduced the time needed for mapping without additional parallelization (i.e., 24 cores x 1 process/core). For 36 total cores, including additional parallelization (12 cores x 3 processes, 16 cores x 2 processes) resulted in an average reduction of about 2 hours in total mapping time. At 44 total cores, we observed slight improvements from additional parallelization (4 cores x 11 processes) relative to the benchmark run lacking parallelization (44 cores x 1 process/core), however, two replicates of the benchmark run required more than 200 GB of memory (kept consistent across all runs) and resulted in out-of-memory (OOM) errors. Finally, for 72 total cores, all configurations of parallel processes yielded similar improvements in efficiency and consistently completed mapping in under 5 hours. Together these benchmarking results show that QTLretrievR is able to scale eQTL mapping efficiently in terms of both memory usage and total runtime. We recommend choosing a minimum number of cores such that dividing the total available cores by this value leaves a remainder of about 4. Then select a total core count that maximizes the number of parallel processes (e.g. 68 cores: 8 cores x 8 processes, 52 cores: 12 cores x 4 processes).

We applied the same overarching parallelization techniques to our mediation analysis step. The primary difference between mapping and mediation is that *r/qtl2*(1) already includes parallelization to speed up processing, whereas *r/intermediate*(4) does not. The inclusion of extra parallelization methods in QTLretrievR maximizes computational efficiency and overcomes the limitations of core saturation in *r/qtl2* and lack of parallelization in *r/intermediate*.

### 3.2. LOD Thresholding

Significance thresholds for QTL analyses are most commonly established empirically through permutation testing. For each permutation, phenotype values are randomly shuffled among individuals, QTL mapping is performed on the shuffled data, and the maximum LOD score from the permuted data is recorded. This process is repeated hundreds to thousands of times, an extreme value distribution is fit to the permuted maximum LOD scores, and in most cases the score corresponding to an alpha of 0.05 is set as the cutoff for calling a significant QTL. For molecular QTL datasets containing thousands of phenotypes, performing a large number of permutations on every phenotype becomes computationally prohibitive. QTLretrievR follows published molecular QTL studies(4–6, 10–13) and performs a rank-based inverse normal transformation (aka “Rank Z transformation”) on normalized phenotype values prior to QTL mapping and permutation testing(20). This data transformation step serves two purposes: first, it eliminates the need for non-parametric marker regression models that are generally less powerful than parametric mapping models; and second, it makes the distribution of permutation LOD scores more comparable across phenotypes in a large dataset. As a result, a significance threshold derived from a subset of Rank Z-transformed phenotypes should be representative of the full phenotype set. To estimate the numbers of subsetted phenotypes and permutations that maximize statistical rigor while minimizing computational expense, we took advantage of a recently published eQTL analysis in mouse neural progenitor cells (NPCs) where we performed permutation testing individually for every expressed gene (“gene-specific thresholding”; n = 14163 genes total)(6). For this analysis, we considered the results of gene-specific thresholding (n = 4116 significant eQTL) as our ground truth set. Next, we evaluated how different combinations of phenotype subsets and permutation counts affect equivalence and accuracy to identify generalizable LOD threshold settings that minimize computational cost while controlling Type I and Type II errors. We performed permutations ranging from 500 - 1000 on a random subset of autosomal genes (the phenotype of interest in eQTL mapping) ranging from 10 - 100. For each subset, we calculated the median significant LOD score among the subsetted genes and compared the resulting significant/non-significant eQTL calls to the ground truth classifications using gene-specific thresholding (Figure 1). Our calculations showed that all combinations of gene numbers and permutations were equivalent to using individual gene thresholds on average (TOST equivalence test, epsilon = 0.03), however that same metric was only met 96% of the time when sampling 50 genes with 625 permutations per gene. Although threshold accuracy is improved by increasing the number of subsetted genes, increasing the number of permutations per gene above 500 provides diminishing returns. Using our published eQTL dataset, we identified a “sweet spot” of 75 genes and 750 permutations per gene that provides equivalent results to gene-specific thresholding (>99%) with fewer total permutations. We therefore established the default permutation settings for LOD thresholding in QTLretrievR to randomly select 100 autosomal genes/phenotypes and perform 750 permutations per gene/phenotype.

**Figure 1.**
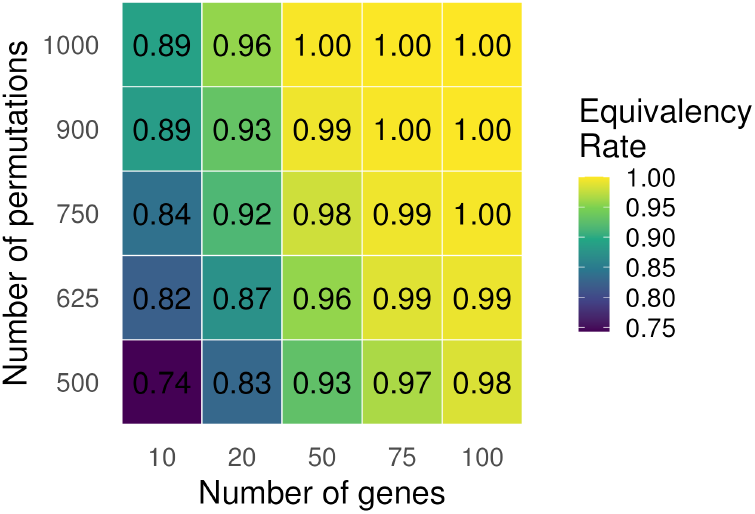
Median LOD significance thresholds derived from permutation testing on random subsets of autosomal genes closely mirror ground truth thresholds derived from gene-specific permutations. Heatmap showing the frequency of equivalence outcomes (TOST, ϵ = 0.03) across combinations of permutation counts and gene numbers. Colors represent the proportion of permutations classified as equivalent.

### 3.3. Downstream Analysis and Visualization

QTLretrievR performs more than molecular QTL mapping, significance testing, and initial analyses of individual peaks. To demonstrate its capabilities on -omics scale data, we revisited the Islet eQTL dataset that was used for benchmarking the mapping step(11). Of note, this islet dataset is part of a larger multi-tissue, multi-omic study on a large number of DO mice(11, 17–19). QTLretrievR is designed to handle data from multiple tissues; mapping results from individual tissues are returned within the same object, thereby simplifying downstream analyses and visualization.

When performing molecular QTL analysis, one of the first requested visualizations is an eQTL map that relates the genome location of a molecular phenotype (e.g. expressed gene, region of open chromatin) to the location associated with its variation (e.g. eQTL peak location). The provided plotting function can work with annotated peaks from multiple tissues and allows for the customization of the plot per tissue. In the mouse islet eQTL map (Figure 2a), we observe the hallmark diagonal band observed in most eQTL studies and indicative of abundant local eQTL. Indeed, the most significant eQTL typically map close to the gene they regulate, and likely reflect variants that act in cis, for example by disrupting a TF binding site in an enhancer region. Points off the diagonal are eQTL that map further apart from their target gene. These distant eQTL exert trans effects on target gene expression usually through an intermediate, termed a “mediator” gene. These long-range intra- or inter-chromosomal effects are likely conferred via direct, proximal (cis) effects on the expression or function of the mediator gene, which then influences the expression ofthe distant eQTL target gene(s). Further, we observe vertical bands in the eQTL map indicative of many distant eQTL colocalizing together to form eQTL “hotspots” or “trans-bands”. We can use the ‘transbands’ function in QTLretrievR to identify the chromosomal locations of these hotspots, their boundaries, and how many suggestive and significant QTLs they contain, along with a histogram showing their relative locations and sizes (Figure 2b). eQTL hotspots like the ones seen in Figure 2b are of particular interest because they likely contain important trans-regulatory genes with broad effects on the expression of many target genes.

**Figure 2.**
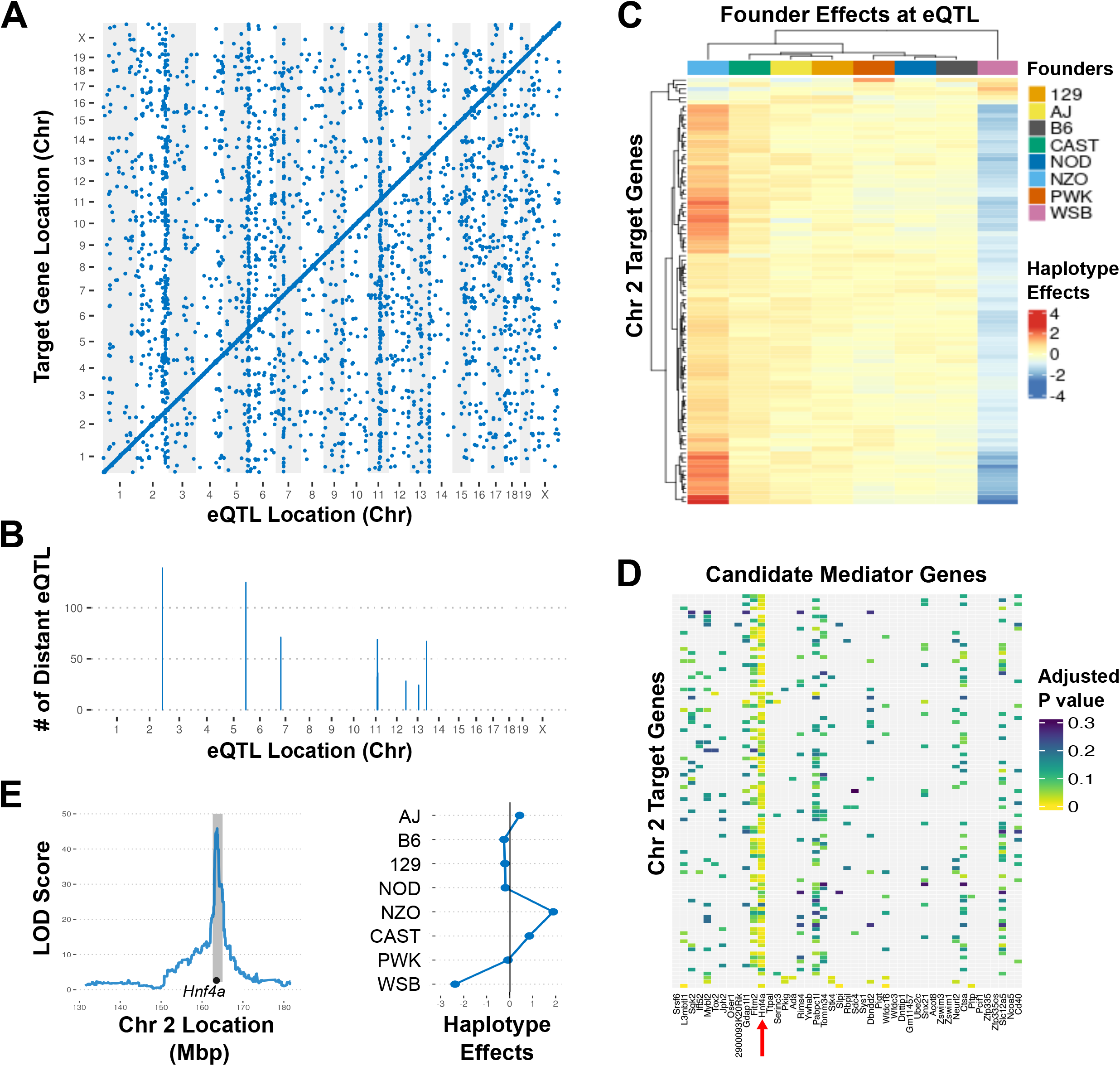
Example visualizations produced by QTLretrievR for islet transcriptomic eQTL analysis. A) An eQTL map plotting the genomic locations of significant eQTL on the x-axis and their target genes on the y-axis. B) Histogram showing the number of distant eQTLs clustering into transbands. C) Heatmap representing founder allele effects for the targets of the Chr 2 hotspot (LOD >9). D) Heatmap showing the mediation LOD drop for candidate mediators of the Chr 2 hotspot (LOD >9). Rows represent hotspot eQTL target genes, and columns show the candidate mediator genes in the Chr 2 region. *Hnf4a* is highlighted with a red arrow. E) Peak plot and founder allele effects for the candidate mediator for the Chr 2 hotspot.

Focusing on individual hotspots, we can further analyze them by looking at the founder haplotype effects of all the target eQTL and identify candidate mediators by integrating mediation results. QTLretrievR has built in functions that collate and plot the target QTL haplotype effects and integrate mediation results with target eQTL. Visualizing the founder haplotype effects at a hotspot allows us to identify groups of target genes whose expression variation is likely driven by one or more founder strain alleles at that locus. For example, the Chr 2 eQTL hotspot is driven by NZO/HlLtJ (NZO) and WSB/EiJ (WSB); DO mice that inherited the NZO founder allele in this region show higher expression and those that inherited the WSB allele show lower expression, respectively, for most target genes (Figure 2c). This haplotype pattern is informative and indicates that the genotype of the causal variant driving this hotspot is likely to vary between NZO and WSB. Further, we can identify a candidate mediator by integrating the mediation analysis results and the trans-band analysis results. The results from this integration are returned as a plot showing the magnitude of LOD score drop from mediation with each candidate gene in the hotspot locus, and the best candidate mediator can be identified by visual inspection. For example, mediation of the Chr 2 hotspot with *Hnf4a* transcript abundance leads to the most significant LOD drop in most target eQTL (Figure 2d), and *Hnf4a* has a local eQTL with similar genotype effects to those observed for the target genes (Figure 2e)—recapitulating the initial study showing this transcription factor as the top candidate mediator for this eQTL hotspot(11). Indeed, Keller and colleagues went on to show that HNF4A was the most significantly enriched transcription factor binding motif in the cis regulatory regions of Chr 2 hotspot target genes, underscoring the power of this systems genetics approach for inferring biologically-relevant gene interactions.

In closing, we’ve designed QTLretrievR to lower the barriers to systems genetic analyses by scaling permutation testing for large multi-tissue and multi-omic studies, integrating molecular QTL mapping and downstream mediation analyses, and providing standardized analysis pipelines and publication-ready plotting functions. Future improvements to the package will include added functions for network inference from the eQTL and mediation results, variant prioritization for individual eQTL, and power analysis for molecular QTL mapping studies.

## Acknowledgments

We would like to acknowledge QTLretrievR beta testers including Catherine Brunton, Madison Armstrong, and Courtney Willey. We would also like to thank Alexa Michaels, Jessica Choi, and Yehya Barakat for their feedback on the manuscript.

## Funding

This work was supported by National Institutes of Health grants GM133495 and R24OD030037 to S.C.M.

## Data Availability

QTLretrievR source code is available at https://github.com/deweyhannah/QTLretrievR. A comprehensive vignette can be found at https://deweyhannah.github.io/QTLretrievR/articles/QTLretrievR.html. The pancreatic islet transcriptomics dataset is available at https://churchilllab.jax.org/qtlviewer/attie/DO500HFD. The DO NPC RNA-seq dataset and associated objects for eQTL mapping can be downloaded from figshare at http://doi.org/10.6084/m9.figshare.28265894.

## Conflict of Interests Statement

The authors declare no conflicts of interest.

## Notes

https://churchilllab.jax.org/qtlviewer/attie/DO500HFD

https://figshare.com/articles/dataset/Cross_cell-type_systems_genetics_reveals_the_influence_of_eQTL_at_multiple_points_in_the_developmental_trajectory_of_mouse_neural_progenitor_cells/28265894

### Competing Interest Statement

The authors have declared no competing interest.

